# The Arabidopsis Calcium Sensor Calmodulin-like 38 Regulates Stress Granule Autophagy and Dynamics during Low Oxygen Stress and Re-aeration Recovery

**DOI:** 10.1101/2021.01.10.426134

**Authors:** Sterling Field, Whitney Gulledge, Daniel M. Roberts

## Abstract

In response to the energy crisis resulting from submergence stress and hypoxia, Arabidopsis limits non-essential mRNAs translation, and accumulate cytosolic stress granules (SG). SGs are phase-separated mRNA-protein particles that partition transcripts for various fates: storage, degradation, or return to translation after stress alleviation. Here, it is shown that RNA stress granules are dynamically regulated during hypoxia stress and aerobic recovery via two phases of autophagy that require the AAA^+^ ATPase CDC48 and the calcium sensor Calmodulin-like 38 (CML38). CML38 is a core hypoxia response-protein that associates with hypoxia-induced SGs. We show that CML38 is essential for SG autophagy during extended hypoxia. Further, cml38 mutants show disorganized SG morphology during extended hypoxia, suggesting a role in SG formation and maintenance. We also show that upon the return of aerobic conditions, intracellular calcium and CML38 are necessary for SG breakdown and turnover, and for upregulating autophagy. *cml38* mutants not only lose these responses, but also have aberrant, sustained autophagosome accumulation during the reoxygenation recovery phase. The findings suggest that CDC48 RNA granule autophagy (“granulophagy”) is conserved in plants, and that the hypoxia-induced calcium sensor CML38 regulates SG autophagy during anaerobic stress as well as during the reprogramming phase associated with reoxygenation.

## INTRODUCTION

Cytosolic mRNA ribostasis is a dynamic process through which the cell partitions selective mRNAs between translating polysomes and non-translating membrane-less mRNP granules in response to changing environmental and developmental cues (Buchan et al., 2013; Chantarachot and Bailey-Serres, 2018; Riggs et al., 2020). During situations of severe stress, translational repression induces declines in polysome loading that is coordinated with the redistribution of transcripts as arrested pre-initiation complexes to liquid-liquid phase separated mRNP assemblies known as stress granules (SGs) (Protter and Parker, 2016). SGs are postulated to serve as “RNA triage centers”, that serve as sites of mRNA storage and partitioning between translating mRNPs (polysomes) and sites of mRNA degradation (P-bodies) (Kedersha et al., 2005; Anderson and Kedersha, 2008; Decker and Parker, 2012; Chantarachot and Bailey-Serres, 2018; Riggs et al., 2020). Such coordinate regulation of mRNA translation, stability, and localization helps to manage limited cellular resources and fine tune gene expression, as well as prepare the cell for recovery programs upon the alleviation of stress conditions.

An example of exquisite cytosolic ribostatic control from higher plants is the reshuffling and reprograming of mRNA translation caused by flooding and submergence stress (Branco-Price et al., 2008; Mustroph et al., 2009; Juntawong et al., 2014; Lee and Bailey-Serres, 2021). Limiting oxygen induced by submergence leads to a decline in respiration and an energy crisis that triggers a variety of adaptive responses to conserve energy and adjust to the hypoxic state (Bailey-Serres et al., 2012; Voesenek and Bailey-Serres, 2015; Cho et al., 2021). Among the responses in the model plant *Arabidopsis thaliana* are a coordinated change in gene expression and mRNA translation to limit protein synthesis from transcripts encoded by non-essential genes and prioritize the expression of small core collection of “hypoxia response proteins” (Branco-Price et al., 2008; Mustroph et al., 2009; Lee and Bailey-Serres, 2021). Decreases in polysome loading induced by hypoxia simultaneously results in a redistribution of translationally-arrested mRNAs to SGs from which the core hypoxia response mRNAs are excluded (Sorenson and Bailey-Serres, 2014). Upon reoxygenation (recovery) the transcription and translation of the core hypoxia transcripts is suppressed (Branco-Price et al., 2008), and stress granules dissipate, releasing mRNA back to translating polysomes (Sorenson and Bailey-Serres, 2014). Ribostasis of cytosolic mRNA by selective translation, storage, and degradation, help cells regulate mRNA and protein synthesis during hypoxia and recovery.

Another factor that needs to be considered in cellular ribostasis is the role of autophagy, a conserved eukaryotic process that allows selective degradation and recycling of cellular material during stress (Yang and Bassham, 2015; Marshall et al., 2019). In Arabidopsis, extended periods of submergence lead to the expression of autophagy related genes (ATGs) and induction of autophagic flux (Chen et al., 2015). While hypoxia-induced autophagy is essential for optimal survival to low oxygen stress (Chen et al., 2015), less is known regarding the specific autophagic targets during extended hypoxia, and additionally, during the process of reoxygenation recovery. The importance of selective autophagy in the homeostatic control of mRNA, mRNP particles, and ribosomes is becoming increasingly recognized (Frankel et al., 2017). Granulophagy is the process of selective SG autophagic clearance in yeast and mammalian systems (Buchan et al., 2013), and defects in this process has been linked to cellular dysfunction that lead to disease states such as Amyotrophic Lateral Sclerosis (Monahan et al., 2016). Autophagic degradation of RNA granules in plants is less studied, but it has been postulated to be a component of plant RNA quality control in response to stress (Hafren et al., 2018; Yoon and Chung, 2019).

In the present paper we show that autophagy is regulated in two phases during hypoxia stress and reoxygenation recovery in Arabidopsis, the first during extended low oxygen stress, and the second a calcium-dependent autophagy burst early in reoxygenation recovery. SG are subject to autophagic turnover during both phases, and consistent with mammalian granulophagy, the ubiquitin segregase CDC48 is required for both. Moreover, we show that *CALMODULIN LIKE 38* (*CML38*), a core hypoxia response gene that encodes a rgsCaM-like calcium sensor that localizes to SG, is required for SG granulophagy, as well as the calcium-dependent reoxygenation autophagy response. Mutants lacking this calcium sensor are defective not only in SG turnover, but show abnormal SG morphology, and a disrupted autophagy program during reoxygenation. Overall, the results show that plant SG are subject to autophagic regulation by a CDC48-dependent mechanism similar to yeast and mammal granulophagy, and that this process is regulated by the CML38 calcium sensor both during extended hypoxia as well as reoxygenation recovery.

## RESULTS

### Autophagy in Arabidopsis roots is stimulated during extended hypoxia and during early reoxygenation recovery

To establish the dynamics of autophagy during hypoxia and reoxygenation recovery, epidermal root cells of 10-day old Arabidopsis plants within the root transition zone were visualized and analyzed as shown in Fig. S1. Visualization of autophagosomes was accomplished by two approaches: 1. Staining of wild type plants with the acid-sensitive fluorescent dye monodansylcadaverine that accumulates within acidified vesicles and has been used to identify autophagosome vesicles (Bassham, 2015); and 2. Live cell imaging of autophagosome formation with the Arabidopsis *GFP-ATG8e* reporter line (Xiong et al., 2007).

Hypoxia results in a redistribution of the MDC stain from a diffuse pattern to readily apparent foci consistent with the appearance of autophagosomes (Fig. 1A). Return to aerobic conditions results in a decrease in MDC-stained foci that are no longer detected by 8 hr after restoration of oxygen (Fig. 1A). Similarly, hypoxia induces the appearance of *GFP-ATG8e* foci consistent with the formation of autophagosomes (Fig. 1B). However, tracking of the GFP-ATG8e signal during reoxygenation showed a complex pattern. Similar to MDC-stained vesicles, return to aerobic conditions results in the decline in GFP-ATG8e autophagosomes during extended recovery (Fig. 1B and C). However, *GFP-ATG8e* plants show a surprising burst of autophagy immediately upon transfer from hypoxic conditions to air, with the number of GFP-ATG8E foci increasing several-fold by 10 min after oxygenation (Fig 1D). The data show that autophagy is part of the response of the plant to sustained low oxygen stress, similar to previous observations with submergence (Chen et al., 2015), but also that autophagy is acutely and transiently induced during reoxygenation recovery.

**Fig 1.**
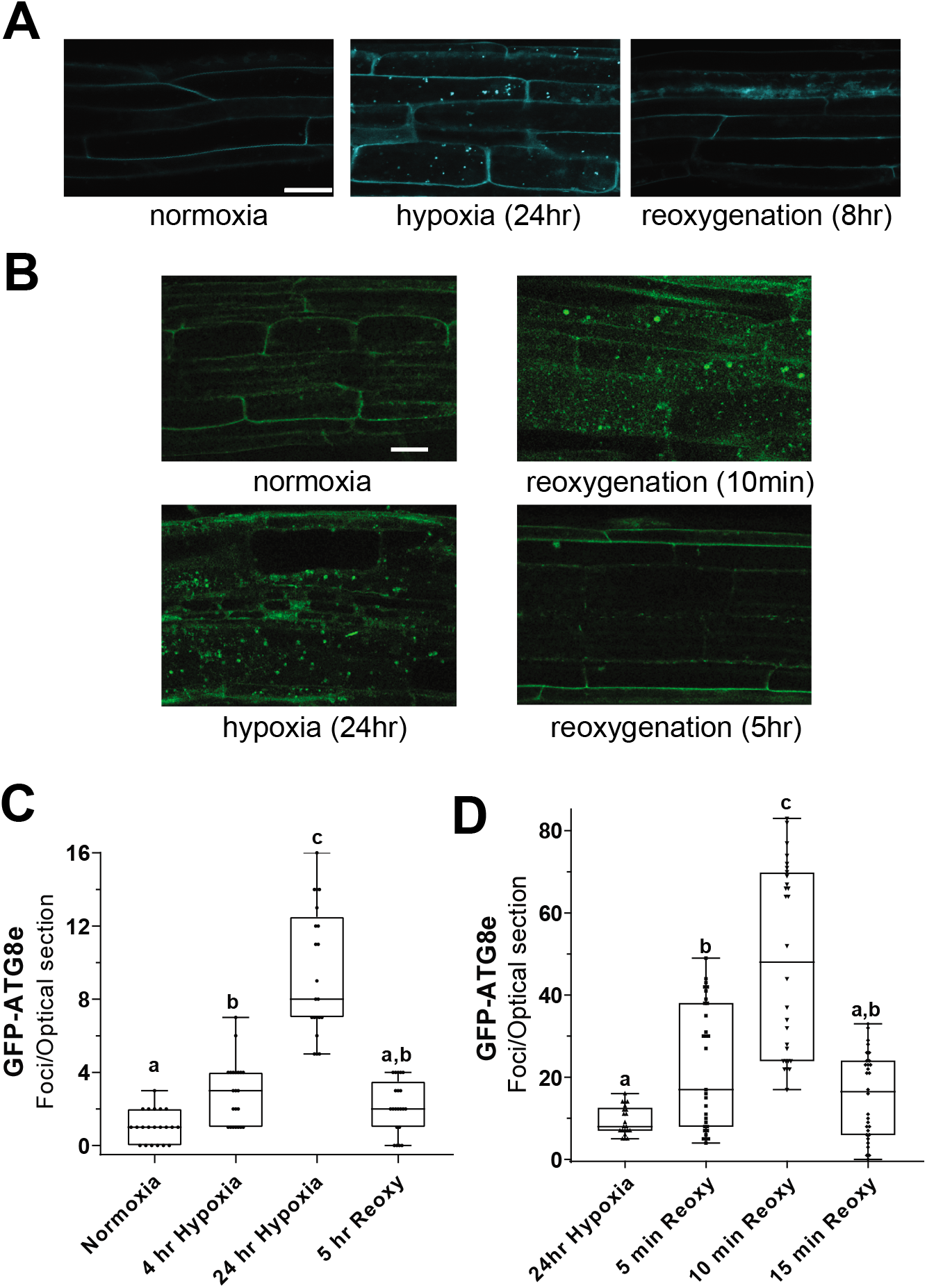
Autophagy is induced by extended hypoxia and shows a burst of activity during early reoxygenation. **A.** Confocal micrographs of root epidermal cells of MDC-stained, 10 day old Arabidopsis Col-0 (wild type) seedlings subjected to argon gas-induced hypoxia followed by reoxygenation recovery. **B.** Confocal micrographs of 10 day old *GFP-ATG8e* seedlings subjected to hypoxia at the indicated times followed by return to aerobic conditions at 24 hr. Scale bars are 25 μm. **C.** Quantitation of GFP-ATG8e foci under the conditions shown in Panel B. **D.** Rapid induction of GFP-ATG8e foci during early reoxygenation. Each data point in panels C and D was obtained from a 2 μm optical section through a single cell. The horizontal line indicates the position of the median value. Different letters represent statistically significant differences (p-value <0.05) based on a One-way ANOVA test.

### RBP47B stress granules are degraded by a CDC48 granulophagy-like process during extended hypoxia

To assess the formation of SG and their potential regulation by autophagy during hypoxia and recovery, transgenic plants with the stress granule marker *RBP47B-CFP* were analyzed. RBP47B is a triple RNA recognition motif protein that can be used to track SG formation and dynamics during cellular stress (Weber et al., 2008). Upon hypoxia treatment, the RBP47B-CFP signal redistributes to foci characteristic of cytosolic stress granules (Fig. 2A). Upon return to aerobic conditions, the numbers of RBP47B foci substantially increase (on average three-fold) in the short term (1 hr) before eventually dissipating by 8 hours of recovery (Fig. 2B). This early reoxygenation phenomenon will be discussed further in a section below.

**Fig. 2.**
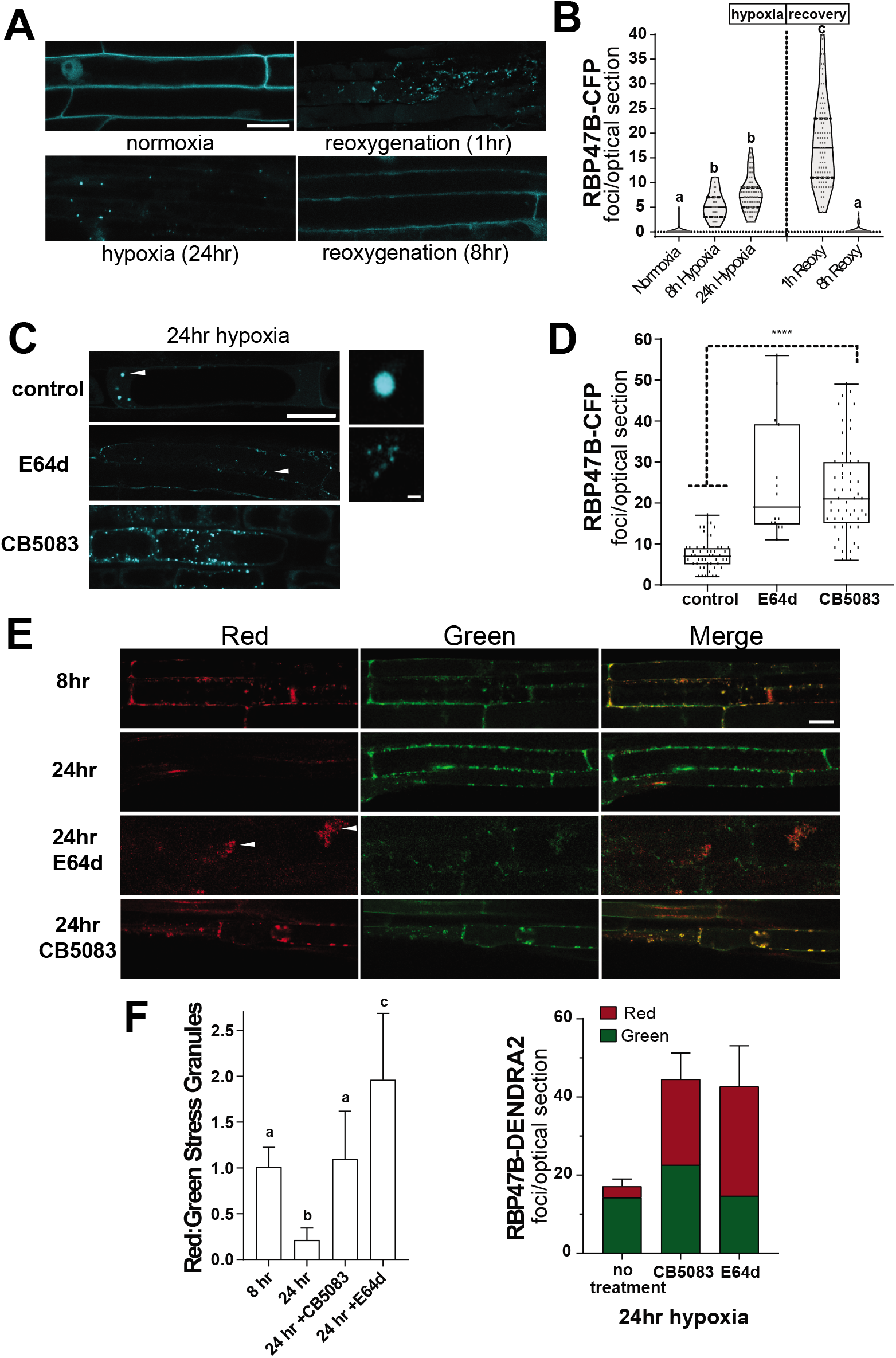
RBP47B granules are degraded by CDC48-dependent autophagy during extended hypoxia. **A.** Confocal micrograph of representative root epidermal cells from *RBP47B-CFP* plants during a hypoxia and recovery time course. **B.** Violin plot showing the quantitation of RBP47 foci at various times of hypoxia and reoxygenation. The solid line indicates the median and the dotted lines indicate the 1st and 3rd quartile values. **C.** Confocal micrograph of root epidermal cells from 24 hr hypoxia-stressed *RBP47B-CFP* plants treated as follows: **Control**, no treatment; **E64d**, in the presence of 1 μM E64d; **CB5083**, in the presence of 2 μM CB5083. A high magnification of the RBP47B particles indicated by the arrowheads is shown to the right. **D.** Quantitation of RBP47-CFP foci from 24 hr hypoxia-stressed *RBP47B-CFP* plants treated as in panel C. **E.** Confocal micrographs of epidermal root cells from *RBP47B-DENDRA2* plants photoconverted prior to hypoxia by the protocol in Fig. S2. Images show the RBP47B-DENDRA2 foci obtained with green or red filters. E64d and CB5083 treatments were done as in panel C. **F.** *Left plot*, the ratio of red to green foci; *right plot*, the actual red to green foci numbers in each cell section under the indicated treatment. Scale bars are 25 μm on panels A, C, and E, or 1 μm on panel C close up. Each data point in panels B, D, and F represents the value from a single cell optical section. Statistical significance was assessed by One way ANOVA analysis, with different letters in panels B and F representing statistical significance (p<0.05).

To investigate whether stress granules are subject to autophagic degradation, the effects of the autophagy inhibitor E64d on the accumulation of RBP47B foci was evaluated (Fig. 2C and D). E64d is a cysteine protease inhibitor that blocks autophagy at the stage of vacuolar degradation (Bassham, 2015; Yang and Bassham, 2015; Marshall and Vierstra, 2018). In Arabidopsis roots this leads to the accumulation of undegraded autophagic body aggregates in the vacuolar compartment (Merkulova et al., 2014). E64d treatment of *RBP47B-CFP* plants resulted in a three-fold increase in the average number of hypoxia-induced RBP47B-CFP SG compared to untreated controls. (Fig. 2D). Confocal microscopy shows that RBP47B-CFP foci accumulate in clusters (Fig. 2C) similar to previous observations of E64d-induced autophagic bodies in Arabidopsis (Merkulova et al., 2014; Bassham, 2015; Marshall and Vierstra, 2018; Marshall et al., 2019).

In mammalian and yeast systems, autophagic turnover of RNA granules requires AAA^+^-ATPases of the CDC48/VCP-p97 ubiquitin segregase family (Buchan et al., 2013; Frankel et al., 2017). CB5083 is an inhibitor that specifically binds to the ATPase domain of CDC48 and inhibits its remodeling function (Marshall et al., 2019; Tang et al., 2019). During hypoxia stress, *RBP47B-CFP* plants accumulate three-fold higher levels of SG in CB5083-treated roots compared to untreated controls (Fig. 2C and D). This observation, together with the results with E64d, supports the hypothesis that SG are subject to CDC48-dependent autophagy during hypoxia. However, it does not exclude the possibility that CDC48 inhibition causes greater production RBP47B foci simply through the induction of new SG aggregates rather than the inhibition of SG autophagy. To resolve these possible interpretations, the temporal dynamics of SG accumulation were tracked with a photo-convertible RBP47B-DENDRA2 as a fluorescent SG marker (Fig. 2E).

The strategy for RBP47B-DENDRA2 (Fig. S2) involves photoconversion from a green to a red fluorescent form by a brief UV pulse before the initiation of the hypoxia stress. The red (old RBP47B) and green (new RBP47B) signal is tracked during the hypoxia time course. Over the first 8 hr of hypoxia, equivalent numbers of SG with red and green RBP47B-DENDRA2 signal accumulate (green/red ratio = 1, Fig. 2F) with substantial co-localization of the two signals (Mander’s co-localization co-efficients, MC_red:green_ and MC_green:red_ = 0.5 – 0.6, Table 1). However, at 24 hr hypoxia the majority of SG contain green RBP47B (a 5-fold higher ratio of green/red SGs, Fig. 2F) while the red RBP47B signal is greatly reduced (Fig 2F). This is also reflected in co-localization analysis that shows that most green SG do not co-localize with red SG (MC_green:red_ =0.04 Table 1). This observation shows the selective loss of SG from early hypoxia (marked by red RBP47B) during extended hypoxia.

**Table 1.**
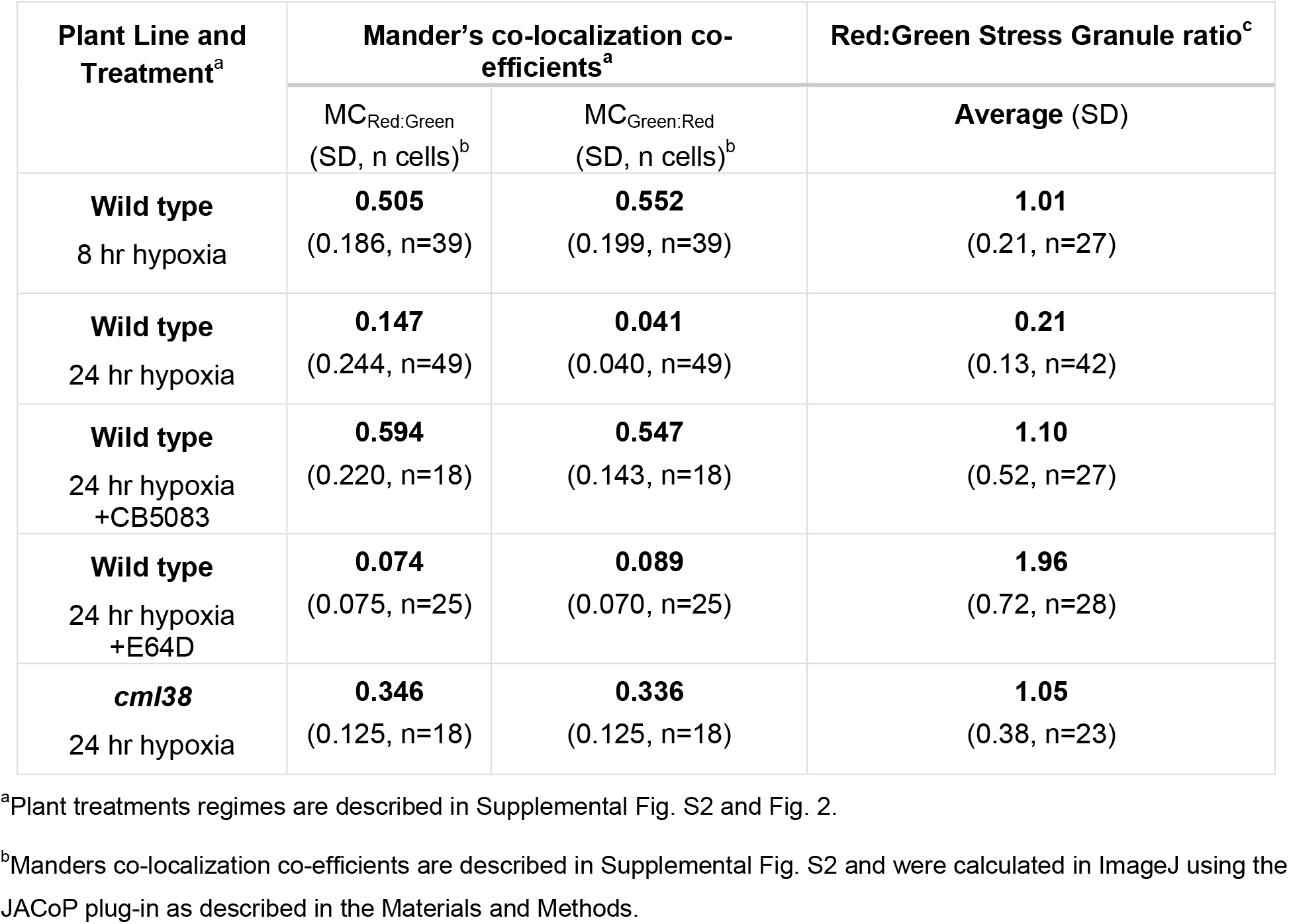
Co-localization co-efficients from *RBP47B-DENDRA2* Experiments

To determine whether this loss is the result of CDC48-dependent autophagy, the DENDRA2 experiment was repeated in the presence of E64d and CB5083 (Fig. 2E, F). In the presence of CB5083, *RBP47B-DENDRA2* plants show the persistence of red RBP47B SG at 24 hr hypoxia, with the numbers of red and green RBP47B SG nearly equal (Fig. 2E and F) and strong co-localization of the two signals (MC_red:green_ and MC_green:red_ = 0.55 – 0.6, Table 1). This observation argues that the elevation of RBP47B SG by the inhibition of CDC48 (Fig. 2D, F) is the result of an inability to turnover SGs during hypoxia rather than from the increased production of new stress granules.

Treatment with E64d shows the accumulation of higher quantities of red RBP47B particles compared to control plants (Fig. 2F). Moreover, the red RBP47B granules preferentially accumulate within aggregated clusters characteristic of autophagic bodies (Fig. 2E), while most green SG are found dispersed outside of these aggregates (Fig. 2E; MC_red:green_ = 0.074, Table 1). Taken together, the data suggest that during sustained hypoxia SG are turned over through autophagy that requires the ATPase activity of CDC48, and that a mechanism similar to yeast and mammalian SG granulophagy is conserved in plants.

### The core hypoxia protein Calmodulin-like 38 shows autophagic turnover during extended hypoxia

CML38 is a core-hypoxia response protein and a calcium-sensor EF hand protein that is acutely induced during hypoxia stress and accumulates within cytosolic granules including SG (Lokdarshi et al., 2016). A hypoxia/reoxygenation recovery time course of *CML38-3YFP* recombineering plants was conducted to compare the accumulation CML38 granules and their potential turnover by autophagy (Fig. 3). The CML38-3YFP signal is not detected under normal oxygen conditions (Fig. 3), but is induced during hypoxia with the fluorescent signal accumulating within cytosolic foci at 8 hr (Fig. 3A, B) consistent with the appearance of SG as previously described (Lokdarshi et al., 2016). However, under sustained hypoxia (24 hr), CML38 foci are no longer apparent in most cells (Fig. 3A, B). Upon return to aerobic conditions, the number of CML38 cytosolic foci increase substantially during the first hour of recovery (Fig. 3 A,B), similar to observations with RBP47B SG (Fig. 2). However, by 8 hr of reoxygenation recovery the CML38 fluorescent signal and overall CML38 protein levels decline and are no longer detected (Fig. 3).

**Fig. 3.**
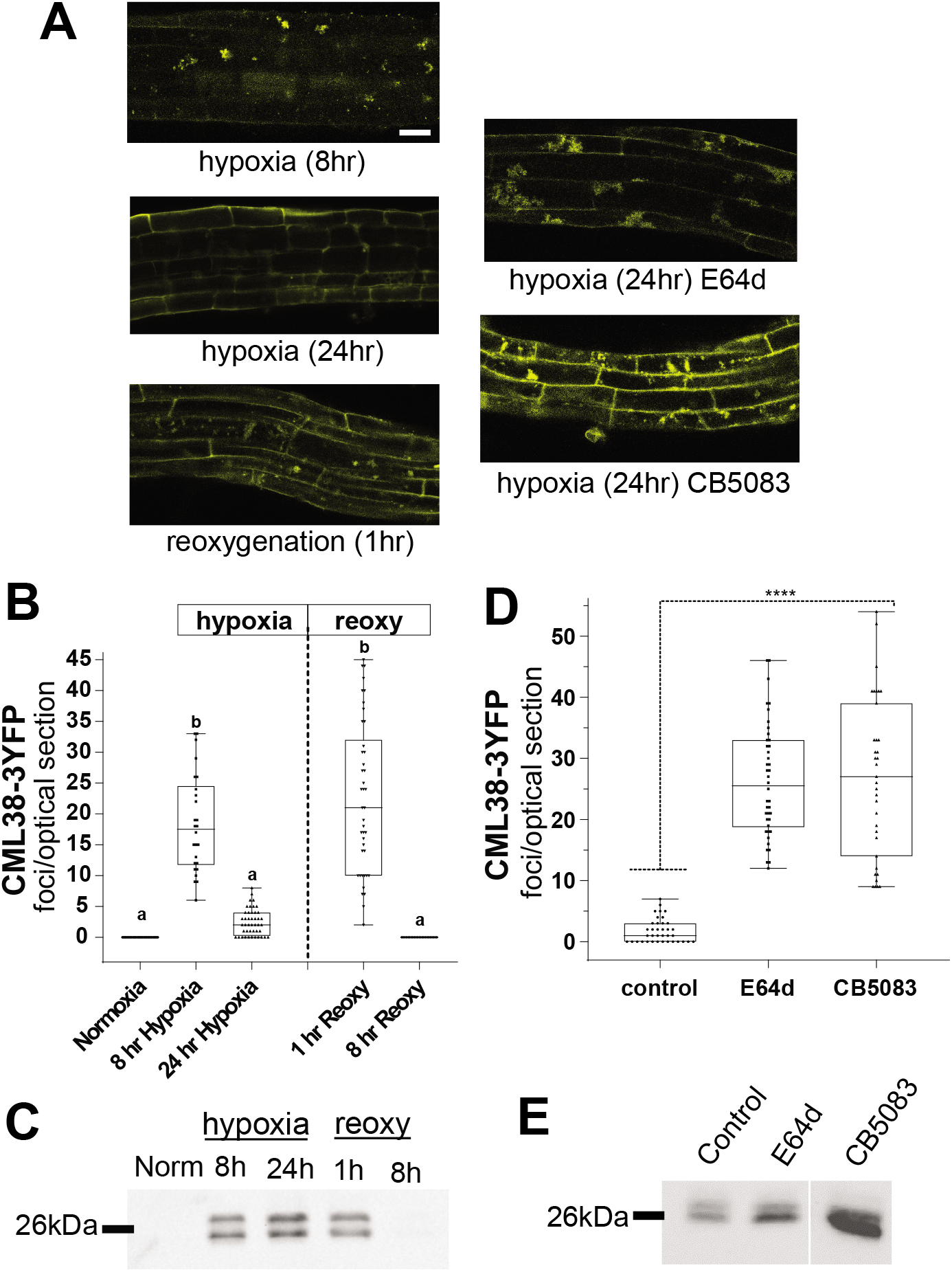
CML38 granules show hypoxia and recovery dynamics similar to RBP47B SG and are regulated by CDC48-dependent autophagy. **A.** *Left*, representative confocal micrographs of root epidermal cells from 10 day old Arabidopsis *CML38-3YFP* recombineering seedlings subjected to hypoxia and reoxygenation recovery as in Figure 2. *Right*, root cells from plants subjected to 24 hypoxia in the presence or absence of 1 μM E64d and 2 μM CB5083. Scale bar is 25 μm. **B.** Quantitation of CML38 foci from *CML38-3YFP* recombineering seedlings plants during a hypoxia and reoxygenation time course. **C.** Western blot comparison of CML38-flg appearance during a hypoxia time course and recovery of 10 day old *CML38p:CML38-flg* plants. **D.** Quantitation of CML38 foci from *CML38-3YFP* plants treated with E64d or CB5083 as described in panel A. **E.** Western blot comparison of *CML38-flg* protein from seedlings of *CML38p:CML38-flg* plants subjected to 24 hr hypoxia with the indicated treatments. Each lane in panels C and E represents 40 μg of total seedling protein. For plots in panels B and D, each data point represents a value from an optical section through a single cell. Statistical significance was determined by One way ANOVA.

To determine whether CML38 foci are subject to CDC48-dependent autophagy, the effects of E64d and CB5083 were assessed (Fig. 3D, E). Treatment of *CML38-3YFP* plants with E64d results in the accumulation of CML38-3YFP signal in granule-like structures that accumulate in clusters (Fig. 3D), similar to the autophagic body-like structures observed with RBP47B stress granules (Fig. 2). Treatment with CB5083 also increases the fluorescent signal with greater numbers of granule-like foci apparent (Fig 3D,E). These observations suggest that autophagy via a CDC48-mediated mechanism may be responsible for the degradation of CML38-3YFP granules during extended hypoxia.

To test this further, the levels of CML38 protein during hypoxia were analyzed by using *CML38-flg* plants. *CML38-flg* plants contain a transgene encoding CML38 with a carboxyl-terminal flag-tag driven by the native *CML38* promoter (Fig. S3). Analysis of the CML38 protein levels show that the protein accumulates to high levels during hypoxia, remains high at 1 hr reoxygenation, and then declines and disappears during sustained reoxygenation (Fig. 3C). Treatment of *CML38-flg* plants with E64d and CB5083 increased the levels of CML38-flg protein at 24 hr hypoxia (Fig. 3E). Taken together, the present data, combined with previous observations that CML38 co-localizes with SG (Lokdarshi et al., 2016), argue that CML38 accumulates in hypoxia-induced stress granules that are degraded via a CDC48-dependent autophagy process.

### Perturbation of CML38 expression alters SG autophagy and morphology during extended hypoxia

The CML38-homolog from tobacco, rgsCaM, is postulated to stimulate autophagy during virus infection (Nakahara et al., 2012; Li et al., 2017). Given its similarity to rgsCaM, and the fact that it localizes to SG, the involvement of CML38 in SG autophagy during hypoxia was explored. To track the appearance and dissipation of SG during hypoxia and recovery, *RBP47B-CFP* reporter plants in wild type and *cml38* mutant backgrounds were compared (Fig. 4). In both wild type and *cml38* backgrounds, RBP47B-CFP SG are absent under normoxic conditions, accumulate in response to hypoxia, and dissipate upon reoxygenation recovery (Fig. 4A). WT and *cml38* plants also accumulate equivalent quantities of cytosolic RBP47B foci with no discernable difference in size or morphology during the first eight hours of hypoxia (Fig. 4A, C). However, during extended hypoxia (24 hr) *cml38* plants accumulate substantially greater quantities of a RBP47-CFP SG foci compared to wild type plants (Fig. 4A). In addition, *cml38* RBP47B SGs at this time point show abnormal morphology compared to wild type plants (Fig. 4C). Quantitation of SG particle shape revealed that *cml38* SGs were significantly less circular and showed irregular shapes compared to wild type SG (Table 2).

**Fig. 4.**
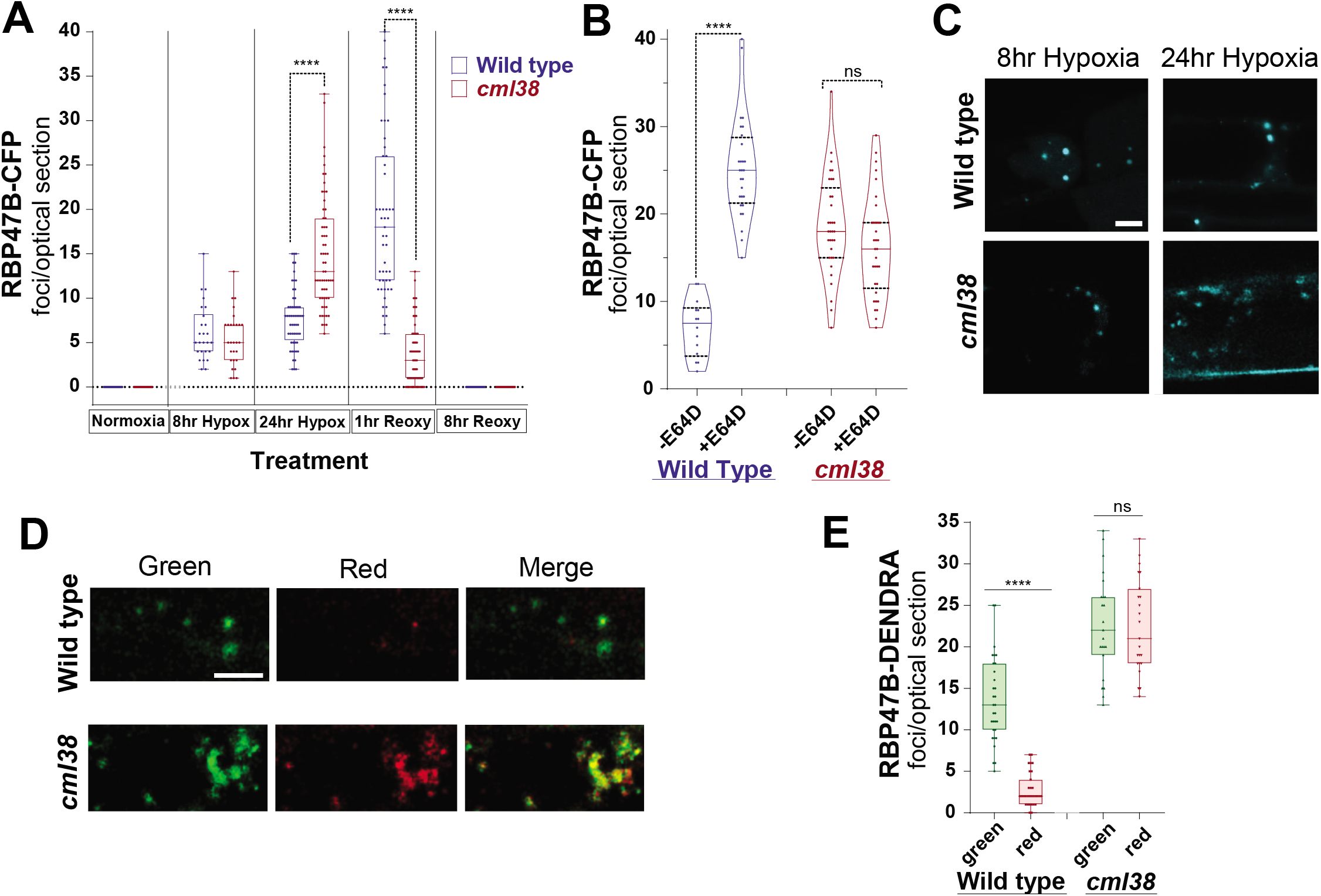
SG autophagy and morphology are disrupted in *cml38* mutant plants. **A.** Quantitation of RBP47B foci during a hypoxia time course and subsequent reoxygenation in *RBP47-CFP* reporter plants constructed in wild type and *cml38* backgrounds. Statistically significant differences between wild type and *cml38* values at each treatment are indicated. **B.** Comparison of the RBP47B foci in epidermal root cell after 24 hr hypoxia in *RBP47-CFP* wild type and *RBP47-CFP cml38* plants in the presence or absence of the autophagy inhibitor E64d. **C.** Confocal micrographs of representative RBP47B-CFP SG foci in WT and *cml38* mutant plants at 8 hr and 24 hr hypoxia. Scale bar is 5 μm. **D.** Comparison of green vs red RBP47B-DENDRA2 foci in root epidermal cells of *RBP47B-DENDRA2* reporter plants in wild type and *cml38* backgrounds after 24 hr hypoxia. Scale bar is 5 μm. MCred:green = 0.04 for the wild type image; MCred:green = 0.37 for the *cml38* image. **E.** Quantitation of red and green RBP47B-DENDRA2 foci in wild type and *cml38* mutants at 24 hr hypoxia. Statistical significance was assessed by One way ANOVA analysis.

**Table 2.**
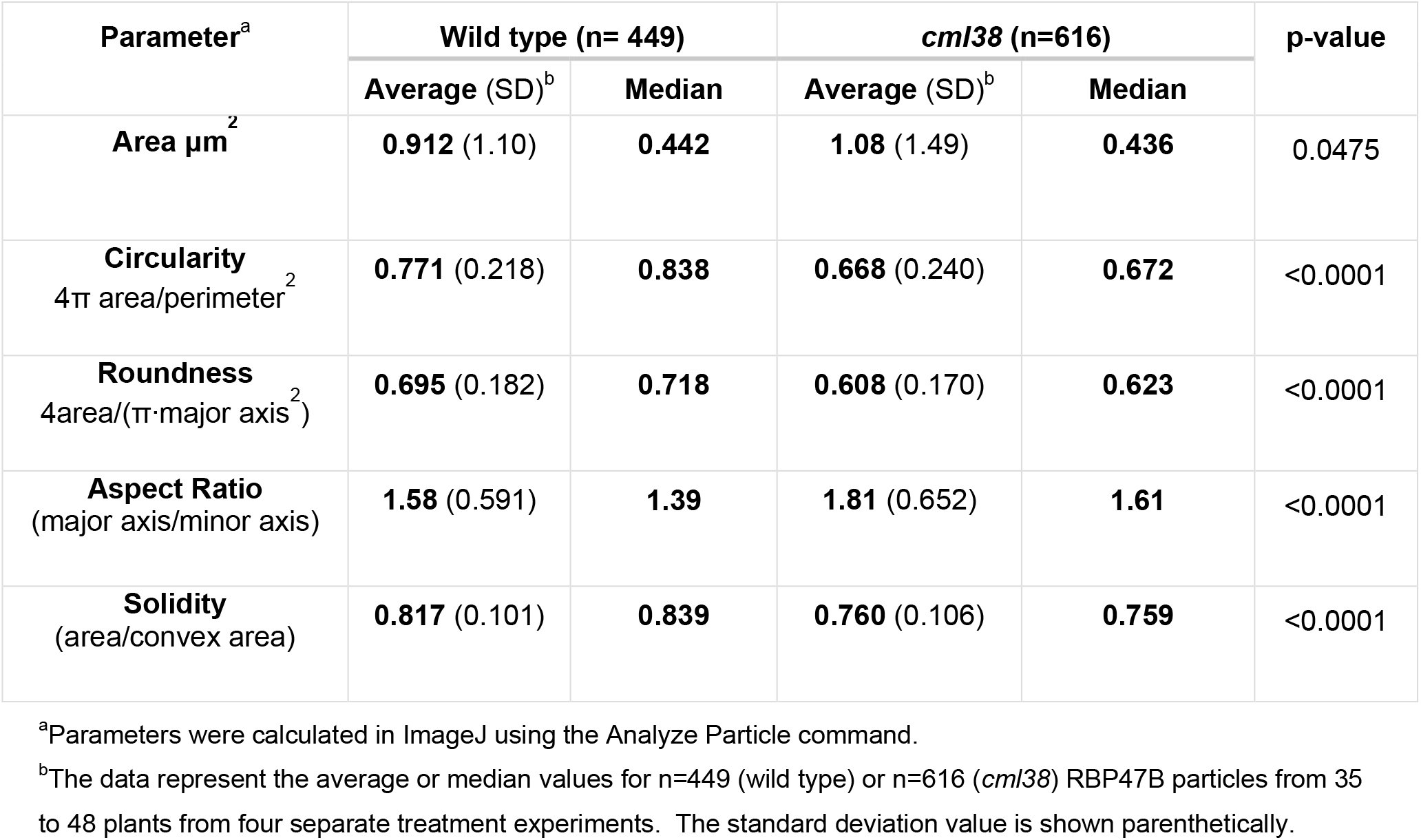
Comparison of RBP47B SG morphology between wild type and *cml38* plants at 24 hr hypoxia

The increased accumulation of RBP47B-CFP SG in *cml38* plants could reflect either increased SG production or inhibition of SG degradation (e.g., by autophagy). To resolve these possibilities, the effect of the autophagy inhibitor E64d on RBP47B foci levels was investigated in wild type and *cml38* RBP47B-CFP reporter plants. Similar to previous observations (Fig. 3), E64d increased the numbers of RBP47B foci in wild type plants. In contrast, *cml38* plants showed no difference in the numbers of RBP47B foci upon E64d treatment (Fig. 4B). This observation suggests that *cml38* plants accumulate greater numbers of stress granules due to the inability to degrade these structures by autophagy.

To investigate this further, *cml38* plants containing a *RBP47B-DENDRA2* transgene reporter were generated and analyzed. As noted above, wild type plants showed a selective loss of the red RBP47B signal during sustained hypoxia (Fig. 4D). However, under the same conditions, *cml38* plants accumulate higher levels of both green and red RBP47B granules in an equivalent ratio (RBP47B_green_/RBP47B_red_ = 1.05, Table 1). Moreover, co-localization analyses reveal a significantly higher co-localization of the red and green SG signal in *cml38* plants (MC_green:red_ = 0.34) compared to wild type (MC_green:red_ = 0.04), (Table 1). This shows that RBP47B SG are degraded to a higher extent over time in wild type, compared to *cml38* plants. Similar to the findings with RBP47B-CFP, *cml38* RBP47B-DENDRA2 granules also show aberrant morphology at 24 hr hypoxia (Fig. 4D). Overall, the data suggest that CML38 is necessary for the autophagic degradation of SGs during hypoxia, and the maintenance of SG structure.

### Perturbation of CML38 expression and CDC48 inhibition alters calcium-dependent SG fractionation during reoxygenation

The number of RBP47B SG foci increases substantially during the first hour of reoxygenation before eventually being cleared as recovery proceeds (Fig. 2). Analysis of the size distribution of RBP47B granules during the first hr of reoxygenation recovery shows a decrease in mean size of the foci (Fig. S4). Further, the mobility of the RBP47B structures increases during this reoxygenation period and larger RBP47B granules dissociate into smaller less organized structures (Supplemental Movies 1-3). These observations suggest that SG are fractionated into smaller aggregates early in the reoxygenation recovery response.

One of the effects of reoxygenation after hypoxia in Arabidopsis roots is an increase in intracellular calcium (Sedbrook et al., 1996). To examine the potential influence of calcium on SG dynamics during reoxygenation, *RBP47B-CFP* plants were imaged in the presence of the Ca^2+^ ionophore ionomycin and either Ca^2+^ or the chelator EGTA (Fig. 5). Ca/ionomycin (Ca Iono) did not affect SG fractionation, with the sizes and numbers of particles showing no difference from 1 hr reoxygenation controls (Fig. 5B, Fig. S4). However, substitution of EGTA for calcium (EGTA Iono) resulted in the loss of the effects of reoxygenation on SG dynamics, with the numbers and size distributions of SGs showing no statistical difference from 24 hr hypoxia controls (Fig. 5B, Fig. S4). This observation suggests that increases in intracellular calcium may be required for the initiation of SG disassembly during early reoxygenation. In support of this, transfer of EGTA-ionomycin treated seedlings back to media containing calcium (EGTA Iono/Ca) triggers SG fractionation within 20 min of treatment (Fig. 5B).

**Fig. 5.**
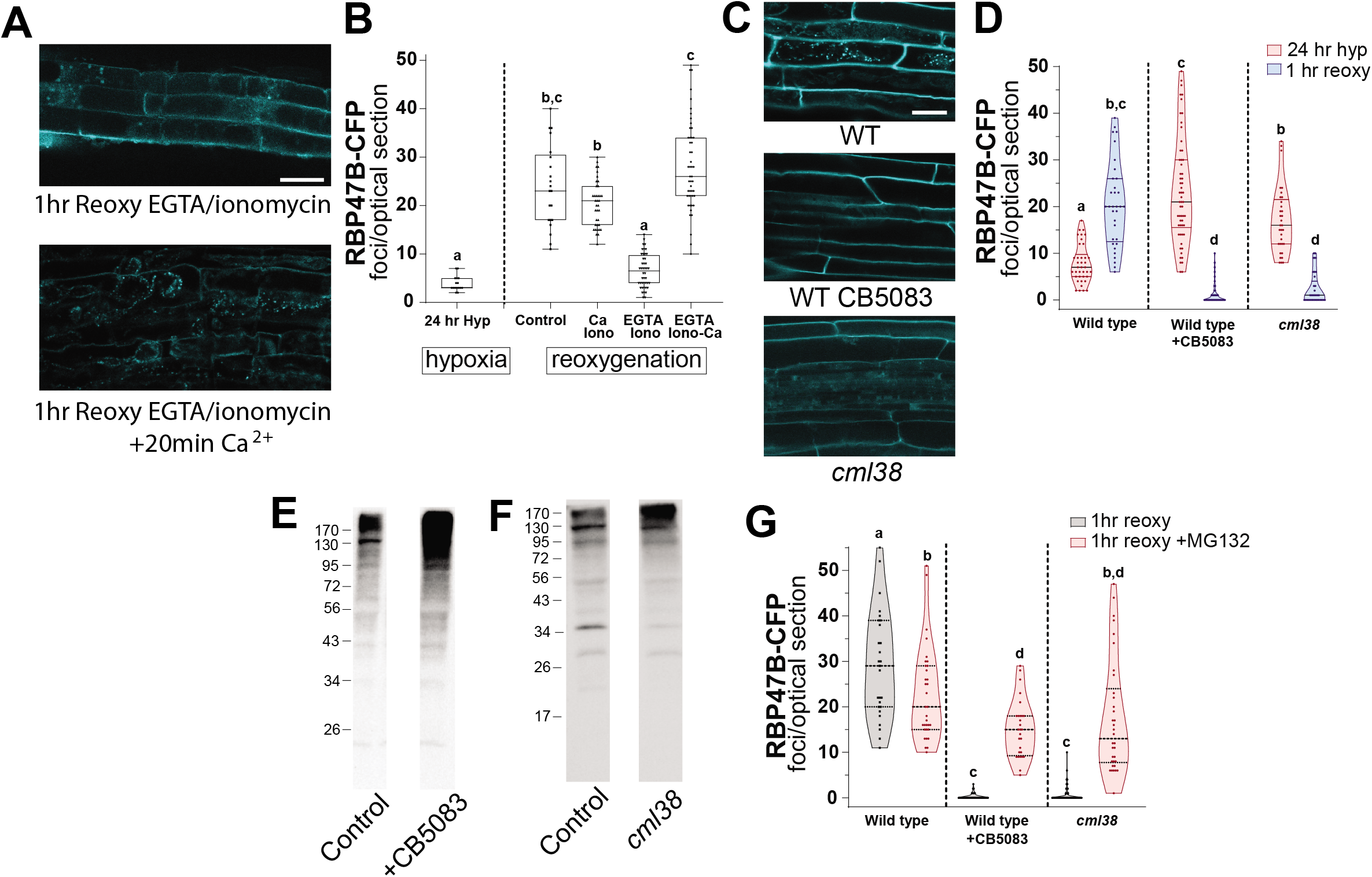
RBP47B granule fractionation into smaller aggregates during reoxygenation requires calcium and CML38. **A.** Representative confocal image of epidermal root cells from *RBP47B-CFP* plants subjected to 24 hr hypoxia followed by 1 hr reoxygenation in the presence of 10 μM ionomycin and 5 mM EGTA (*top*) and 20 min after return to 10 μM ionomycin and 5 mM CaCl_2_ (*bottom*). **B.** RBP47B foci in **RBP47B-CFP** plants subjected to 24 hr hypoxia and 1 hr after return to aerobic conditions. **Control**, no ionomycin; **Ca iono**, 10 μM ionomycin + 5 mM CaCl_2_; **EGTA iono**, 10 μM ionomycin + 5 mM EGTA; **EGTA iono-Ca**, EGTA iono conditions followed by transfer to 10 μM ionomycin + 5 mM CaCl_2_ for 20 min. **C.** Representative confocal image of epidermal root cells subjected to 24 hr hypoxia and then 1 hr reoxygenation from the following plants/conditions: **WT**, wild type *RBP47B-CFP* plants; **WT CB5083**, wild type *RBP47B-CFP* plants + CB5083 (2μM); ***cml38*** and *RBP47B-CFP* plants in the *cml38* background. Scale bar is 25 μm in panels A and C. **D.** Quantitation of RBP47B-CFP foci from plants treated as described in panel C. **E.** Anti-ubiquitin western blot of wild type plants subjected to 24 hr hypoxia in the presence (+CB5083) or absence (control) of 2 μM CB5083. **F.** Anti-ubiquitin western blot of wild type and *cml38* plants subjected to 24 hr hypoxia. Each lane in panels E and F contain 35 μg of protein from whole seedling extracts. **G.** Comparison of RBP47B-CFP foci in root epidermal cells from plants treated as in panel C, with (red plots) and without (gray plots) 50 μM MG132. Each data point in panels D and G represents the RBP47B foci from optical sections of a single cell with the median and quartile positions shown on a violin plot. Statistical significance was assessed by One way ANOVA analysis.

Since the CML38 calcium sensor is localized to SG-like structures, the possibility that the calcium-dependent SG fractionation during reoxygenation requires CML38 was investigated with *RBP47B-CFP cml38* plants (Fig. 5D). While extended hypoxia results in the accumulation of greater numbers of RBP47B-CFP SG in *cml38* plants compared to wild type, 1 hr reoxygenation did not show the reoxygenation increase in RBP47B particles observed in wild type plants. Rather, surprisingly, *cml38* plants showed a near complete loss of RBP47B foci within 1 hr of return to reoxygenation (Fig. 5D). If CML38 is required for autophagic clearance of SG, why would *cml38* mutants exhibit increased loss of SG during reoxygenation?

One potential answer comes from the cross-talk between the proteasome and autophagy pathways as homeostatic mechanisms for the clearance and degradation of cellular components. Inhibition of one pathway can lead to compensation by the upregulation of the other (Kocaturk and Gozuacik, 2018). In the case of SG clearance during reoxygenation, this may be the case (Fig. 5). For example, the inhibition of CDC48 by CB5083 leads to elevation of SG (Fig. 2) and an increase in the levels of ubiquitinated proteins (Fig. 5E) during extended hypoxia. However, plants treated with CB5083 show a rapid and complete loss of SG upon reoxygenation (Fig. 5D) that is blocked by the proteasome inhibitor MG132 (Fig. 5G). This suggests that inhibition of CDC48 granulophagy leads to the degradation of SG components by the proteasome pathway during reoxygenation. Examination of *cml38* mutants show a similar phenomenon. The levels of ubiquitinated proteins are elevated in *cml38* mutants during hypoxia (Fig. 5F), and the rapid loss of SG in the absence of CML38 is also blocked by MG132 (Fig. 5G). The data suggest that CML38 and calcium may be required as part of the dynamic remodeling and clearance of SG during reoxygenation in wild type plants, and that the proteasome is responsible for SG degradation when autophagy is impaired during recovery.

### Sustained autophagy in *cml38* plants during reoxygenation recovery

The finding that *cml38* mutant plants are defective in SG autophagy lead us to evaluate whether the autophagy process itself is defective in *cml38* mutants (Fig. 6). To compare the dynamics of autophagosome biogenesis and disappearance, transgenic *YFP-ATG8e* reporter lines were constructed in wild type (Col-0) and *cml38* mutant backgrounds. The accumulation of MDC stained vesicles and *YFP-ATG8e* autophagosomes in wild type and *cml38* plants occurs in a nearly indistinguishable fashion upon the administration of hypoxia (Fig. 6 A-D). Thus, despite the observation that *cml38* mutants show defects in the selective turnover of SG during sustained hypoxia, *cml38* plants still exhibit the same pattern of global autophagy induction during low oxygen stress as wild type plants. However, *cml38* mutant plants show substantial differences in autophagy upon reoxygenation. In contrast to wild type plants, which show loss of MDC vesicles and return to a normal state upon reoxygenation, *cml38* plants are insensitive to reoxygenation and maintain a sustained elevated level of autophagosomes even 8 hr after restoration of an aerobic state (Fig. 6A, B). Further comparison of *YFP-ATG8e cml38* reporter lines with wild type show a similar difference, with YFP-ATG8e autophagosomes persisting for hours after reoxygenation in the *cml38* background (Fig. 6C, D).

**Fig. 6.**
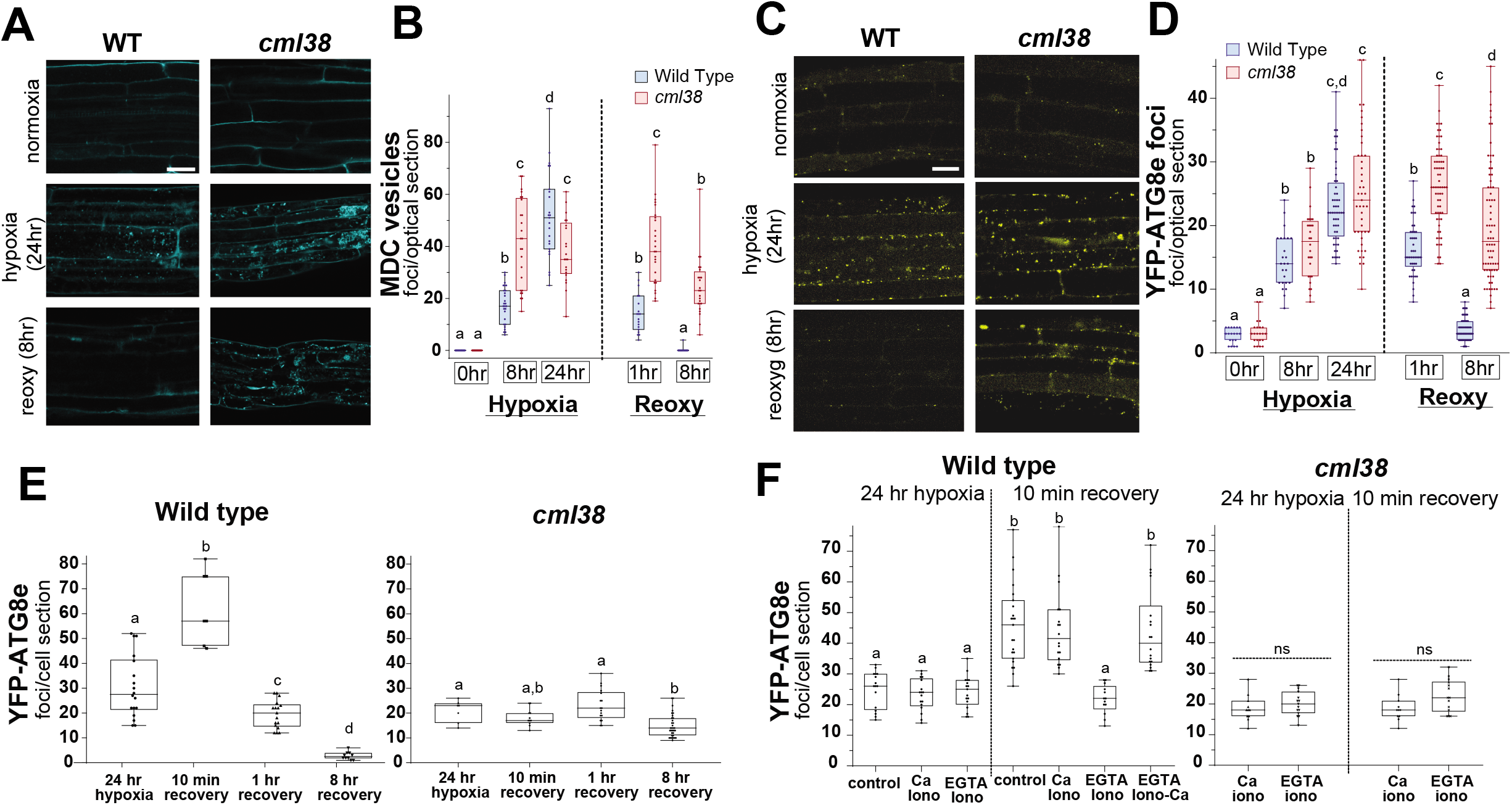
Autophagy regulation during reoxygenation is disrupted in *cml38* mutant plants. **A.** Confocal micrographs of root epidermal cells of MDC-stained wild type and *cml38* plants during hypoxia and reoxygenation time course. **B.** Histogram comparing MDC foci numbers in cell optical sections under the indicated treatments. **C.** Confocal micrographs of YFP-ATG8e foci in root epidermal cells of *YFP-ATG8e* reporter plant lines in wild type and *cml38* backgrounds. Scale bars is 25 μm in panels A and C. **D.** Histogram comparing the numbers of YFP-ATG8e foci in cell optical sections under the indicated treatments. **E.** Analysis of YFP-ATG8e foci during short term reoxygenation in wild type (*left*) and *cml38* (*right*) plants. **F.** *YFP-ATG8e* reporter plant lines were exposed to 24 hr hypoxic conditions before return to aerobic conditions (recovery) in the presence of 10 μM ionomycin and calcium (5 mM CaCl_2_) or the calcium-chelator EGTA (5 mM). **Control**, plants with no ionomycin treatment; **Ca Iono**, plants treated with ionomycin and CaCl_2_ at 23 hr hypoxia and imaged at the times indicated; **EGTA Iono**, plants treated with ionomycin and EGTA at 23 hr hypoxia and imaged at the times indicated; **EGTA Iono-Ca**, EGTA/Ionomycin plants that were transferred to calcium conditions for an additional 20 min prior to imaging. Statistical significance was assessed by One way ANOVA analysis.

As noted in Fig. 1, reoxygenation results in a rapid burst in the appearance of ATG8e foci in the Arabidopsis *GFP-ATG8e* reporter line of (Xiong et al., 2007). A similar reoxygenation burst is observed in the *YFP-ATG8e* wild type plants generated here (Fig. 6E). Comparison of this reoxygenation phenomena in ionomycin and either calcium or EGTA shows that the reoxygenation burst of ATG8e autophagosomes is suppressed by ionomycin/EGTA, but is restored upon resupplying calcium (Fig. 6F). In contrast to *YFP-ATG8e* wild type plants, *YFP-ATG8e cml38* plants show no response to reoxygenation with ATG8e foci remaining at a static unchanging level compared to hypoxia controls, with no apparent effects of ionomycin, EGTA, or calcium (Fig. 6E, F). The data suggest that the calcium and the calcium sensor protein CML38 are both required for the second phase of autophagy that occurs during reoxygenation.

## DISCUSSION

At the onset of low oxygen stress resulting from flooding, land plants prioritize the expression of hypoxia responsive genes involved in adaptation. This coordinate regulation is complex, and occurs in a spatial and temporally regulated fashion at multiple levels including: hypoxia-responsive gene transcriptional activation, epigenetic regulation by DNA methylation, RNA processing/splicing and nucleocytoplasmic transport, selective mRNA translation of hypoxia-responsive transcripts, and the suppression of translation of unnecessary mRNA and the accumulation of cytosolic stress granules (Branco-Price et al., 2008; Mustroph et al., 2009; Juntawong et al., 2014; Cho et al., 2019; Lee and Bailey-Serres, 2019) (reviewed in Cho et al., 2021; Lee and Bailey-Serres, 2021). Collectively, these programs coordinate the survival response to the energy crisis and toxicity associated with low O_2_ stress, as well as prepare the plant for the restoration of an aerobic state. In the present study, we show that autophagy is associated with both the hypoxia and reoxygenation recovery responses in Arabidopsis, that RNA stress granules appear to be regulated by a CDC48-dependent granulophagy process, and that the core hypoxia calcium-sensor protein CML38 regulates elements of both responses.

### Selective autophagy of stress granules involves a CDC48-dependent mechanism

Similar to responses to a wide variety of biotic and abiotic stresses (Yang and Bassham, 2015; Marshall and Vierstra, 2018; Signorelli et al., 2019), sustained submergence stress induces autophagy in Arabidopsis and is essential for optimal survival to this stress (Chen et al., 2015). While the need for recycling of resources during the severe energy crisis associated with low oxygen stress is clear, less is known regarding the specific targets and cargo subject to autophagy during sustained hypoxia. Autophagy *atg* mutants overaccumulate salicylic acid and ROS, suggesting a balancing role for autophagy in modulating ROS signaling while preventing ROS-mediated oxidative damage (Chen et al., 2015). Arabidopsis S-nitroso-glutathione reductase is also a target for selective autophagy during hypoxia, suggesting a potential role in NO signaling (Zhan et al., 2018). The present work shows that stress granule RNA particles that accumulate early during oxygen deficit are also among the targets for selective autophagy during extended hypoxia, suggesting an additional role in RNA homeostasis.

The regulation of RNA homeostasis by autophagy is an emerging area, with various RNAs and ribonucleoprotein particles such as ribosomes and RNA granules regulated by autophagy (Frankel et al., 2017). In yeast and mammalian systems, a pathway known as “granulophagy” (Zhang et al., 2009; Zhao et al., 2009; Buchan et al., 2013; Frankel et al., 2017) mediates the degradation of cytosolic RNA stress granules, such as stress granules, P-granules, and processing bodies. Granulophagy requires the action of the CDC48/VCP-p97 AAA^+^-ATPase (Buchan et al., 2013). CDC48 serves as an ATP-dependent ubiquitin segregase that recognizes and removes ubiquitinylated proteins from complexes, targeting them for autophagy (Ye et al., 2017), and thus ubiquitin-dependent CDC48 remodeling of RNA granules may precede autophagy. This is supported by the observation that CDC48 depletion results in increased ubiquitinylated proteins and impairs SG regulation in human cell cultures (Seguin et al., 2014). The observation that the inhibition of CDC48 blocks autophagy of SG in hypoxia-stressed Arabidopsis supports the proposal that higher plants possess a similar granulophagy pathway.

How selective autophagy of SG is coordinated with RNA homeostasis remains to be elucidated, but may represent part of a larger network of the gene expression regulatory program. The results obtained with *atg* mutants show that defective autophagy exhibits a complex effect on the levels of hypoxia-response transcripts. For example, the classical hypoxia response transcripts that encode *ADH1* and *PDC1* are elevated in *atg* mutants, whereas ethylene-signaling transcripts such as *EIN2, ERF1*, and *HRE1* are significantly lower in *atg* mutants (Chen et al., 2015). Based on analyses of the transcriptome, translatome, and SG transcript populations (Branco-Price et al., 2008; Mustroph et al., 2009; Juntawong et al., 2014; Sorenson and Bailey-Serres, 2014), the hypoxia mRNA regulatory landscape is complex, and changes spatially and temporally depending upon the needs of the cell and the severity and duration of the energy stress (Cho et al., 2021; Lee and Bailey-Serres, 2021). The finding here that SG are targets of hypoxia-induced autophagy argues that this is part of the larger regulatory program that modulates mRNA homeostasis during long term hypoxia.

### CML38 is required for granulophagy and SG maintenance during hypoxia

The calcium sensor protein CML38 is one of the 49 Arabidopsis core hypoxia response genes (Mustroph et al., 2009) that is acutely induced during hypoxia stress and is required for an optimal response to this stress (Lokdarshi et al., 2016). CML38 accumulates within cytosolic bodies during hypoxia that are proposed to be stress granules based on RBP47B co-localization experiments and co-IP/MS analyses (Lokdarshi et al., 2016). The present work shows hypoxia-induced CML38 granules are also targeted for autophagy during extended hypoxia stress in a CDC48-dependent manner. In support of the hypothesis that CDC48 is necessary for CML38/stress granule autophagy, it is interesting to note that CDC48 is among the proteins identified in previous CML38 IP experiments of hypoxia-stressed Arabidopsis (Lokdarshi et al., 2016).

More importantly, the present work with *cml38* mutants shows that CML38 is required for SG granulophagy during hypoxia. While a mechanism through which CML38 regulates SG autophagy is not yet clear, potential insight comes from work on the homologous “regulator of gene silencing” calmodulin from *Nicotiana*. RgsCaM was initially discovered in tobacco as an interaction target for the potyviral Helper Component Proteinase (Anandalakshmi et al., 2000) and has subsequently been shown to regulate and suppress secondary siRNA silencing pathways by triggering the selective autophagy of components of the RNA silencing machinery (Nakahara et al., 2012; Li et al., 2017; Yang et al., 2019). The amino terminal region of rgsCaM contains an “ATG8 interacting motif” (AIM) that mediates an interaction of rgsCaM with the autophagosome membrane protein ATG8 (Conner et al., 2019). It has been hypothesized that rgsCaM facilitates autophagy by recruiting target proteins to the nascent autophagosome through an ATG8 interaction (Conner et al., 2019). CML38 and other rgsCaM family members have a conserved AIM domain in their amino terminal regions. Whether CML38 regulates autophagy by interacting directly with ATG8, or regulates autophagy in another manner, remains to be elucidated.

In addition to regulation of SG autophagy, *cml38* T-DNA mutants also show defective SG morphology during extended hypoxia. Disruption of normal granulophagy homeostasis in other systems causes similar defects in SG organization and structure. For example, inhibition of granulophagy by RNA_i_ disruption of the CDC48 ortholog VCP in HeLa cells results in abnormal SG morphology and composition (Seguin et al., 2014). *Atg* mutants showed a similar SG phenotype (Seguin et al., 2014), and a role for CDC48/VCP remodeling in SG assembly as well as clearance was proposed (Frankel et al., 2017). The observations with *cml38* mutants suggest a similar function in normal SG assembly and homeostasis.

### Stress granule and autophagy dynamics during reoxygenation recovery

An unexpected finding in the present study is that a second phase of autophagy is induced upon reoxygenation, with autophagosomes appearing and peaking within minutes of the return of molecular oxygen followed by gradual disappearance as recovery proceeds. Recent work suggests that similar rapid and transient autophagic bursts may be a common reprogramming response to several hormonal and environmental signals that helps facilitate the rapid removal of unnecessary cellular components, and is necessary for the establishment of new programs (Rodriguez et al., 2020). In this regard, it important to consider that the post-hypoxia reoxygenation triggers a complex metabolic reprograming response as well as generating its own set of stresses. Reoxygenation restores the expression of transcriptionally and translationally repressed gene products, as well as the concomitant repression of the hypoxia-response genes (Branco-Price et al., 2008). At the same time, the return of aerobic conditions after submergence stress results in the elevation of reactive oxygen species, osmotic stress, and other adverse conditions that trigger a new set of adaptive programs to mitigate oxidative stress (Yeung et al., 2019). The transient induction of autophagy may be a necessary part of a reoxygenation program to re-establish aerobic physiology as well as prevent oxidative and other post-hypoxia damage.

Previous work (Sorenson and Bailey-Serres, 2014) shows that reoxygenation in hypoxia-treated Arabidopsis plants triggers the disassembly of UBP1 SG, and the release of client mRNA back to translating polysomes. In the present study, RBP47B SG also disassemble upon reoxygenation. However, prior to dissipation, RBP47B SG show dynamic structural changes during the first hour of reoxygenation, including fractionation of SG into more numerous and smaller aggregates, and an increase in cellular motility. This behavior is consistent with previous observations of SG disassembly and degradation in mammalian systems (Protter and Parker, 2016; Alberti et al., 2017). SG fractionation and disassembly is an ATP-dependent process that is mediated by the combined activities of RNA helicases, HSP chaperones, and CDC48/VCP AAA^+^-ATPases (Buchan et al., 2013; Walters et al., 2015; Protter and Parker, 2016; Alberti et al., 2017). Recovery from stress involves the fractionation of large mature stress granules by these ATP dependent processes, releasing mRNPs back to translation, and producing smaller aggregates consisting of SG core components that are further disassembled and cleared by autophagy (Wheeler et al., 2016).

The reoxygenation-dependent disassembly of RBP47B SG appears to require the action of CDC48, since the inhibition of this enzyme results in the apparent degradation of SG components through an alternate pathway involving the proteasome. This may be an example of reciprocal regulation of the autophagy and proteasome pathways, which are able to compensate for each other to maintain cellular homeostasis in mammalian systems (Kocaturk and Gozuacik, 2018), including the degradation of cellular aggregates (Minoia et al., 2014) and SG (Turakhiya et al., 2018).

### Calcium and CML38 are necessary for the reoxygenation autophagy and SG responses

Arabidopsis plants with a calcium-reporter aequorin transgene show multiple rapid luminescent transients in root cells upon reoxygenation after extended anoxia (Sedbrook et al., 1996). These signals are blocked by the calcium chelator EGTA, suggesting that they represent intracellular calcium signals associated with early reoxygenation signaling. The present study shows that EGTA blocks both the autophagy burst as well as the early SG fractionation response during early aerobic recovery, and that calcium in the presence of the ionophore ionomycin restores these reoxygenation behaviors. These observations suggest that calcium transients associated with reoxygenation are required for autophagy and SG remodeling during early reoxygenation.

Observations with the *cml38* mutant suggest the calcium sensor protein CML38 is necessary to transduce this calcium signal. While both *cml38* and wild type plants accumulate autophagosomes during hypoxia, *cml38* plants fail to respond to reoxygenation or calcium during recovery. Instead, autophagosomes remain persistent and unchanging for several hours after the return of oxygen. These observations suggest that loss of CML38 could lead to mis-regulation during reoxygenation reprograming that results in constitutive autophagy. A similar constitutive induction of bulk autophagy was observed in Arabidopsis *rns2* mutants with defective selective ribophagy (Hillwig et al., 2011; Floyd et al., 2015).

Mutant *cml38* plants also over accumulate SG during extended hypoxia due to the inability to degrade these bodies via autophagy. However, upon reoxygenation, SG rapidly disappear within one hour. This is similar to the observations with CDC48 inhibition noted above, and decline in SG numbers during *cml38* plant reoxygenation is also blocked by a proteasome inhibitor. This observation suggests that CML38 and CDC48 are not only involved in SG autophagy during extended hypoxia, but could also be necessary for calcium-dependent SG remodeling, autophagy, and clearance during reoxygenation (Fig. 7).

**Fig. 7.**
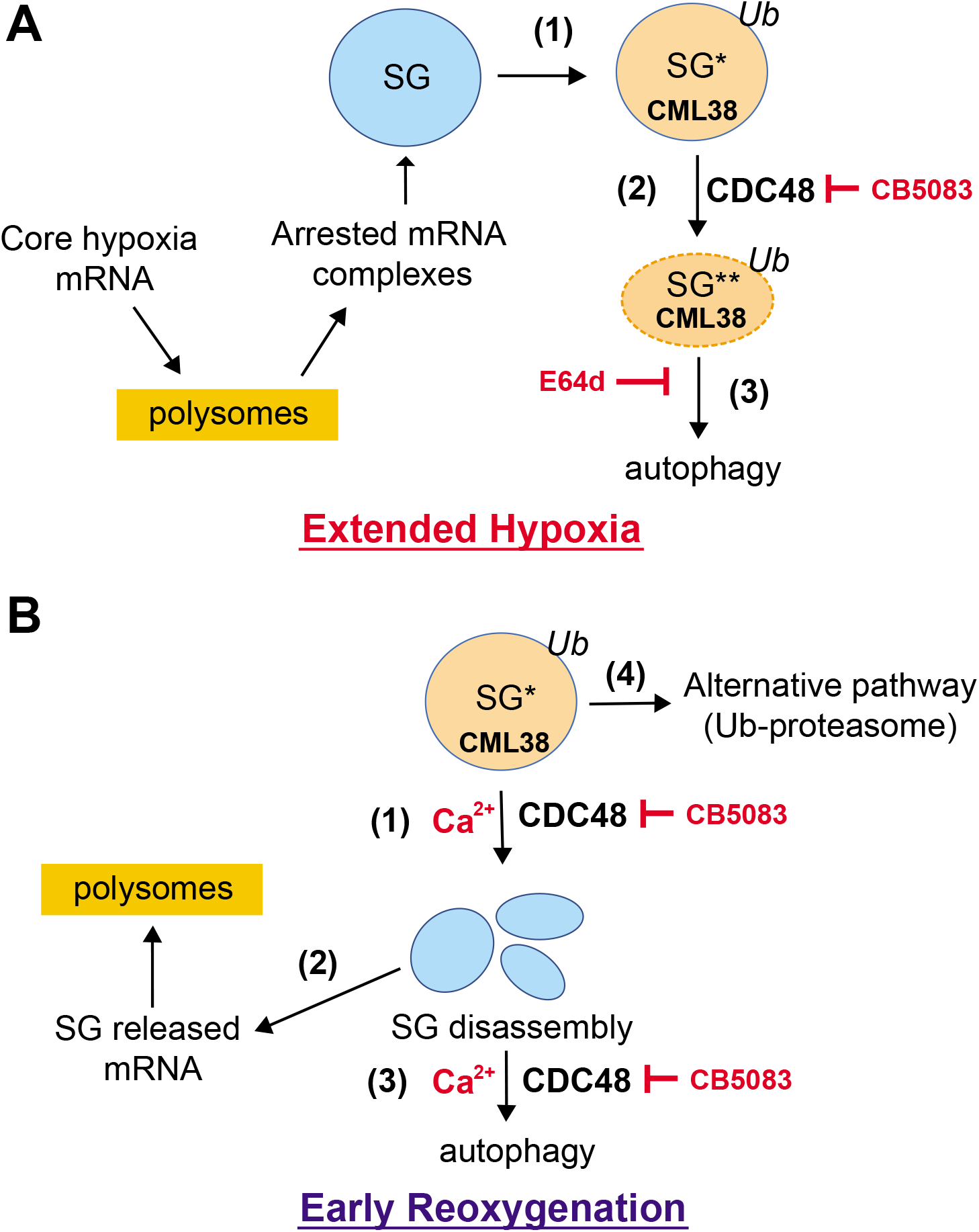
Model for SG regulation by CML38 and CDC48 during hypoxia and reoxygenation recovery. **A.** During extended hypoxia, translationally arrested mRNA complexes accumulate in SG while the core hypoxia response mRNAs are preferentially translated. ***(1)*** Ubiquitinated SG that are targeted for autophagy are bound by CML38. ***(2)*** The ubiquitin segregase CDC48 remodels components of the SG to allow its ***(3)*** degradation through selective autophagy similar to the CDC48/VCP granulophagy processes in yeast and mammalian systems. A subset of SGs that are not targeted for granulophagy (blue) persist. **B. *(1)*** Under the return of aerobic conditions, SG are fractionated into smaller fragments in a calcium, CML38, and CDC48 dependent manner. ***(2)*** Transcripts stored in SGs during hypoxia stress, including those necessary for the recovery response, are released to polysomes for translation. ***(3)*** SG core fragments are degraded by autophagy that is upregulated in a calcium dependent manner during early reoxygenation as part of the reprogramming and recovery response. ***(4)*** Inhibition of this process in *cml38* mutant plants or by inhibition of CDC48 triggers an alternative pathway for SG disassembly that involves the Ub-proteosome. The CML38 calcium sensor appears to be necessary for coordinating the calcium signal that mediates these responses.

## MATERIALS AND METHODS

### Plant Growth and Stress Treatments

All lines used are in the Columbia-0 background. Specific genotypes used in this study are described in the Table S2, and the generation of new lines in this study is described under the Methods. The T-DNA *cml38* mutant plant line is from (Lokdarshi et al., 2016) and the *GFP-ATG8e* autophagosome reporter line is from (Xiong et al., 2007). Seeds were sterilized, plated, and vernalized for 48 hours at 4 **°**C on 1/2x Murashige-Skoog (1/2x MS) media without sucrose, and were grown under long day conditions (16 h light [~100 μmol/m^2^/s]/8 h dark at 22 °C) for 10 days prior to stress treatments. Argon gas hypoxia was administered as described by (Lokdarshi et al., 2016). Hypoxia treatments were conducted with plants at 8 hrs into the day cycle under light conditions (70 μmol/m^2^/s). Hypoxia was administered by purging with Argon gas (AR UHP300, Airgas; <2% O_2_ measured using a Traceable Oxygen probe, Fisher Scientific). Reoxygenation was conducted by return of plates to normoxic conditions in the light (70 μmol/m^2^/s). *RBP47B-DENDRA2* seedlings were photoconverted with a 30 second pulse of UV light administered by a Hoefer Mighty Bright transilluminator prior to hypoxia and reoxygenation treatments.

Autophagy inhibitor treatments were performed by supplementing media with 1 μM E64d as described in (Liu and Bassham, 2010; Huang et al., 2019) prior to treatments. CDC48 inhibitor treatments were performed as described by (Marshall et al., 2019) by supplementing media with 2 μM CB5083. Proteasome inhibitor treatments were performed as described by (Marshall and Vierstra, 2018) by transfer of plants to 1/2x MS liquid media containing 50 μM MG132. To assess calcium dependence, seedlings were incubated in 50 mM Tris-HCl pH 7.5, 150 mM NaCl supplemented with 10 μM ionomycin and either 5 mM EGTA (iono-EGTA) or 5 mM CaCl_2_ (iono-Ca) for one hour prior to imaging. As noted in the text, in some experiments seedlings from iono-EGTA treatments were transferred to iono-Ca conditions for 20 min prior to imaging.

### Molecular Cloning and Transfection Techniques

The *RBP47B-CFP* construct under the control of the Cauliflower mosaic virus (CaMV) 35S promoter was generated in (Lokdarshi et al., 2016). For other constructs, cDNAs containing the open reading frames of *RBP47B, CML38*, and *ATG8e* were generated from PCR of a cDNA template prepared from total RNA from 6 hr hypoxia-treated 10 day old *Arabidopsis thaliana* seedlings. Total RNA was extracted with the E.Z.N.A. Plant RNA Kit (OMEGA) and total cDNA synthesis was performed with the High Capacity cDNA Reverse Transcription Kit with random hexamer primers, according to manufacturer’s protocols (Applied Biosystems). All primers used in this study are in the Table S1.

An amino terminal in-frame translational fusion of *eYFP-ATG8e* was generated in the pEarleyGate 104 plant expression vector system under the control of the CaMV35S promoter (Earley et al., 2006). The *ATG8e* PCR product, which contains attB1 and attB4 sites, was recombined with the pDONR 207 donor plasmid in a BP Clonase II reaction according to manufacturer’s protocols (Invitrogen). *ATG8e* was cloned into pEarleyGate 104 using a LR Clonase II reaction (Invitrogen).

*RBP47B-DENDRA2* was assembled by combining: *DENDRA2* (original plasmid obtained from AddGene) PCR product amplified with primers of overlapping flanking sequence with *RBP47B* and introduced a six glycine linker between the two reading frames; *RBP47B* amplified by PCR using 6 hr hypoxia 10 day old Arabidopsis cDNA; and ligated together into the gateway compatible Gene Entry Clone (GEC) plasmid (Wang et al., 2013) by using Gibson Assembly (New England Biolabs). *RBP47B-DENDRA2* was cloned into pEarleyGate 100 (Earley et al., 2006) under the control of the CaMV35S promoter.

*CML38-flg* driven by the native *CML38* promoter was generated in the promoter-less pEarleyGate 301 vector (Earley et al., 2006). The *CML38-flg* construct was generated with primers that remove the stop codon and introduce an in-frame FLAG tag with a six glycine linker (Fig. S3). *CML38-flg* was assembled by Gibson Assembly by combining the GEC vector (Wang et al., 2013), the *CML38* ORF PCR product, and a 1500 base pair genomic DNA fragment upstream of the *CML38* start site containing the promoter region of the *CML38* gene. The *CML38_ptO_:CML38-flg* construct was then cloned into pEarleyGate 301. All final plasmid constructs were confirmed by Sanger DNA sequence analysis performed on an Applied Biosystems 3730 Genetic Analyzer at the University of Tennessee Genomics Core Facility.

All constructs were transformed into *Agrobacterium tumefaciens* strain GV3101 (Koncz and Schell, 1986) by electroporation and selected on LB-agar with 50 μg/mL gentamycin, kanamycin and rifampicin. *Arabidopsis thaliana* was transformed by the floral dip method (Clough and Bent, 1998), and seeds were selected on ½ MS media with 25 μg/mL glufosinate. *RBP47B-CFP, RBP47B-DENDRA2*, and *YFP-ATG8e* lines were prepared in wild type (Col-0) as well as *cml38* backgrounds. T1 Plants selected on 25 μg/mL glufosinate media were genotyped by amplifying the transgene using PCR on genomic DNA extracted from plant leaves with transgene specific primers. Experiments used one line per construct and were T3 plants homozygous for the transgene.

### Confocal Microscopy Methods

Imaging experiments (Fig. S1) were conducted on Leica SP8 Confocal Microscope system at the Advanced Microscopy and Imaging Center, at the University of Tennessee, Knoxville. Images were captured with a Leica Sp8 laser scanning confocal microscope (Wetzlar, Germany) using a 40x wet mount objective. Root transition zones (Fig S1) were captured (~300 μm x ~150 μm per frame) to include multiple epidermal cells with a z-axis step size of 2 μm for each optical sections (z-series depth of 15-40 μm). The pinhole setting was between 0.8-2 AU for all images. All images were collected using 4x line averaging with bidirectional laser scanning at 600 Hz scanning speed.

For imaging, whole seedlings were mounted in Phosphate Buffer Saline (PBS; 137 mM NaCl, 27 mM KCl, 100 mM Na_2_PO_4_, 18 mM KH_2_PO_4_) on a microscope slide. Fluorescent proteins were detected at (excite/emission): 405 nm/451-550 nm (CFP), 491 nm/501-568 nm (GFP), 514 nm/525-580 nm (YFP), 488 nm/505-530 nm (Green DENDRA2), 543 nm/560-650 nm (Red DENDRA2), 405 nm/475-525 nm (MDC). All images used the HyD hybrid detector. For MDC visualization, seedlings were immersed in 10 μM monodansylcadaverine (MDC) in 1x PBS for 10 minutes before image analysis.

Confocal micrographs were captured and saved using the Leica LASX software Version 3.3.0, and were maintained as .lif files to preserve meta-data until processing and analysis. For processing, all images were first opened in LASX to add scale bars and apply the LASX built-in smooth rendering filter to reduce noise. LASX was used for false coloration, and assembling merged overlays for co-localization analysis.

Confocal micrographs were exported and analyzed in ImageJ (Schneider et al., 2012). Images were smoothed with the ImageJ built-in filter and noise was reduced by background subtraction. Particle quantitation was conducted as described in Fig. S1. Individual cells were outlined and the particles (size distribution 0.1 to 10 μm^2^) were quantitated with the built-in ImageJ Particle Counter (Schneider et al., 2012). Co-localization analysis in *RBP47B-DENDRA2* plants was performed using the Just Another Colocalization Plugin (JACoP) plugin in ImageJ (Bolte and Cordelieres, 2006). Images corresponding to DENDRA2 red and green channels were imported into JACoP and a region of interest was selected that corresponds to the cytosol of an individual cell. Co-localization was quantified by calculation of Mander’s coefficients (Manders et al., 1993). Two Mander’s coefficients were computed: MCC_*red:green*_ quantifies the fraction (0-1) of the red DENDRA2 signal that overlaps green signal, and *MCC_green:red_* quantifies the fraction (0-1) of the green DENDRA2 signal that overlaps red signal. This approach is useful since it takes into account the possibility of disparate co-localization of red and green particles due to unequal quantities of each particle type.

ImageJ was also used to analyze and quantify RBP47B stress granule shape data (Jain et al., 2016). Granules were analyzed by importing tiff files into ImageJ, applying the ImageJ smooth rendering filter and background subtraction to reduce noise, and a threshold was set that highlighted all particles. Granule shape was calculated by the ImageJ built in Particle Analyze command to quantify and output parameter, area, circularity, roundness, aspect ratio, and solidity (Table 2). All microscopy experiments, and all values and data shown in this study were collected from a minimum of three independent biological experiments, with multiple cells from a minimum of three seedlings analyzed for each treatment in each biological replicate. Outliers were assessed by Tukey’s interquartile range analysis. Unless otherwise noted in figure legends, statistical significance (p<0.05) was assessed by multiple comparison analysis using one-way ANOVA with GraphPad Prism 7.00.

### Immunochemical Methods

Ten-day old seedlings were frozen in liquid nitrogen, ground with a mortar and pestle, and resuspended (200 μL buffer for ~0.3 g tissue) for 10 minutes on ice with 50 mM Tris-HCl pH 7.5, 0.15 M NaCl, 10% (v/v) glycerol, 0.01% (v/v) NP-40, 1 mM dithiothreitol, with one Protease Inhibitor Mini Tablet (Thermo Scientific, EDTA free) per 10mL of resuspension buffer. Samples were centrifuged at 4 **°**C for 15 minutes at 16,000 x g, and the supernatant fraction was collected and analyzed by Western blot. Protein concentration of extracts was measured by using a Bradford assay (Bio-Rad). Extract protein (35 to 40 μg) was separated by SDS-PAGE on 12.5% w/v polyacrylamide gels, were electroblotted onto polyvinylidene difluoride (PVDF) membranes, and Western blot analysis and chemiluminescent detection were performed by the method of (Wallace and Roberts, 2005). For CML38-flg immunoblots, a mouse anti-FLAG primary monoclonal antibody was used (anti-DYKDDDDK FG4R, 1 μg/mL overnight at 4 **°**C, Invitrogen), and horseradish peroxidase (HRP)-coupled goat anti-mouse IgG was used as a secondary antibody horseradish peroxidase (0.5 μg/mL for 1 hr at 21 **°**C; Invitrogen). Ubiquitin immunoblotting was performed with a rabbit anti-UBQ polyclonal antibody (0.1 μg/mL for 1 hr at 21 **°**C; AgriSera), and HRP-coupled goat anti-rabbit IgG as a secondary antibody (0.05 μg/mL for 1 hr at 21 **°**C; Novus Bio).

### Quantitative Real Time PCR

Quantitative Real Time PCR (Q-PCR) was done as previously described (Lokdarshi et al., 2016) with modifications. PCR was performed with PerfeCTa SYBR Green SuperMix (Quantabio) using 10 ng of total cDNA per a 12.5 μL reaction. Q-PCR was performed on a BioRad CFX96 Real-Time System with the following parameters: 1 cycle of 5 min at 95 **°**C and 40 cycles of: 1 cycle of 15 sec at 95 **°**C, 1 cycle of 15 sec at 55 **°**C, 1 cycle of 30 sec at 72 **°**C. Quantitation of gene expression was calculated using the comparative threshold cycle (Ct) method as described in (Choi and Roberts, 2007).

### Accession Numbers

Sequence data from this article can be found in the EMBL/GenBank data libraries under the following accession numbers: CML38 (AT1G76650), ATG8e (AT2G45170), RBP47b (AT3G19130).

## Acknowledgements

This work was supported by National Science Foundation Grant MCB-1121465. We thank Dr. Tessa Burch-Smith for guidance and critical comments during preparation of this manuscript. We thank Dr. Diane Bassham, Iowa State University, for providing seeds for the *GFP-ATG8e* line. We also thank Dr. Andreas Nebenfuehr for assistance with confocal microscopy and image analysis. We also thank Dr. Ansul Lokdarshi and Dr. W. Craig Connor for assistance with the generation of RBP47B constructs and transgenic lines as well as advice on image analysis.

## Author Contributions

**S.F.** designed and performed the majority of the experiments, coordinated the contributions of other authors, supervised W.G., and together with D.M.R. wrote the manuscript; **W.G.,** helped to design and performed GFP-ATG8e experiments; **D.M.R**., supervised and conceived the project, provided support and supervision of personnel, was involved in the design of experiments, analysis and interpretation of the data, and co-wrote the manuscript with S.F.

## Supplemental Data

**Figure S1.** Area of Arabidopsis root cell imaging during hypoxia and recovery.

**Figure S2.** Photoconversion and co-localization with RBP47B-DENDRA2.

**Figure S3.** Generation of the *CML38-flg* reporter line.

**Figure S4.** RBP47B granules fractionate in a calcium dependent manner during early reoxygenation.

**Table S1** Oligonucleotides used in this study

**Table S2** Bacteria, plant lines, and plasmid reference table

**Supplemental Movie S1.** Dynamics of SG disassembly during hypoxia in *RBP47B-CFP* plants.

**Supplemental Movie S2.** Dynamics of SG disassembly during early reoxygenation in *RBP47B-CFP* plants.

**Supplemental Movie S3.** SG disassembly during early reoxygenation in *RBP47B-CFP* plants.

